# Bayesian Independent Component Analysis reconstructs independent modules of gene expression

**DOI:** 10.1101/2025.05.22.655463

**Authors:** Jorge Carrasco Muriel, Teddy Groves, Lars Keld Nielsen

## Abstract

Transcriptional regulation—the modulation of gene expression in response to environmental stimuli—is fundamental to cellular function. Identifying groups of co-regulated genes helps elucidate gene functions and characterize how an organism has evolved to respond to various stimuli. In previous works, signal processing algorithms have been applied to characterize the transcriptional regulatory modes, known as iModulons, of bacteria. However, these methods do not quantify uncertainty of the results and are difficult to integrate with different sources of information. In this work, we propose a Bayesian model of Independent Component Analysis that addresses these issues by providing a formal structure to quantify the uncertainty of gene activations and membership of co-regulated genes, achieving state-of-the-art alignment with known regulators. Furthermore, we expand this Bayesian model to explain and integrate first multi-strain and then multi-omics data.

**Author summary:** Understanding how genes are turned on and off is crucial for deciphering how living organisms respond to their environment. Genes often work together in groups, and identifying these co-regulated groups can reveal their functions and how organisms adapt to changes. Previous methods have used complex mathematical techniques to find these gene groups in bacteria, but they come with limitations: they do not measure how confident we can be in the results and are hard to combine with other types of biological information.

In our study, we introduce a new approach using Bayesian statistics to overcome these challenges. This method not only helps us identify groups of co-regulated genes more accurately but also allows us to quantify our confidence in these findings. Additionally, our approach can easily integrate different kinds of data, such as information from various bacterial strains or other biological processes. This makes our method a powerful tool for exploring gene regulation, with potential applications in understanding diseases, developing new treatments and advancing biotechnology.

## Introduction

Transcriptional regulation is central to our understanding of metabolism and cell physiology. Transcription provides the link between genetic information and its mobilization in response to stimuli. Evidence of this linkage has been instrumental to understanding gene function [1], cell development [2] and the molecular mechanisms of disease [3].

Gene regulation is coordinated with genes clustering into modules of expression [4], often mapped to a defined biological function, e.g., general stress response [5]. The evidence of these modules can be statistical, when a group of genes are found to be correlated across experiments or stimuli, or mechanistic, when a known transcriptional factor regulates a group of genes.

The elucidation of mechanistic links between transcription factor binding and gene expression involves expensive and time-consuming experimental methods such as ChipSeq [6]. n contrast, statistical elucidation seeks to deconvolute the modules of transcription directly from transcription data. While sequencing technologies such as RNAseq are streamlined and inexpensive, deconvolution was until recently only modestly successful [4]. This changed with the adoption of signal processing algorithms, which have achieved great success in producing modules of expression—hereafter iModulons—reconstructing with high fidelity the transcriptional regulatory network (TRN) of *Escherichia coli* [7], *Bacillus subtilis* [8] and *Salmonella enterica* [9], among others [10]. Furthermore, iModulons have been applied to successfully transfer cellular functions from organism to organism [11], showing their usefulness as functional components of expression for synthetic biology.

The signal processing algorithm currently used for the disentanglement of iModulons, Independent Component Analysis (ICA), faces two important limitations. First, it lacks uncertainty quantification of the output, leading to complications in model-driven design of experiments. Second, ICA yields a dense matrix that requires post-processing to determine gene membership in each iModulon.

These challenges are particularly acute for non-model or less-researched organisms, where little or no reference information about the regulatory network is available. The lack of a reference TRN in such cases both complicates the post-processing step and limits the biological interpretation of the method, making it difficult to assess whether the available data were sufficient to characterise the latent iModulon regulatory structure. To address these challenges, we introduce Bayesian ICA for gene membership allocation, resolving the postprocessing issue by incorporating membership formally with sparse priors and yielding uncertainty quantification, naturally given by Bayesian probability. The method is evaluated against classical ICA, achieving state-of-the-art alignment to known TRNs in *E. coli*. Additionally, we extend the model to accommodate different data from multiple strains, accounting for general transcription changes in Adaptive Laboratory Evolution (ALE) strains.

Finally, we extend it to include both proteomics and transcriptomics from multiple sources, showing the integration of the different information into shared and refined iModulons.

## Design and implementation

### Data

To aid comparisons with the original iModulon paper [7], we used the PRECISE-1 transcriptomics database of 278 *Escherichia coli* samples hosted at https://github.com/SBRG/precise-db. Reads were transformed to gene log transcripts per millions (TPMs), centered by subtracting the wild-type condition, and whitened by transforming it to a space where the covariance matrix is the identity matrix. We found whitening to be an important step to avoid divergences during gradient-based MCMC sampling.

For the model with multi-omics data, proteomics and transcriptomics were taken from [12] and [13] in the form of number fractions (see Supplementary Material Eq S.16).

### Bayesian model

The Bayesian model of ICA was implemented in the Stan Probabilistic Programming Language version 2.34.0 [14]. MCMC sampling was performed using the No-U-Turn Sampler (NUTS) algorithm [15]. All models discussed in this manuscript yielded no post-warmup divergent samples and did not hit the maximum tree depth. The python preprocessing scripts and Stan code for the models can be found at https://github.com/teddygroves/bicata.

A detailed description of the Bayesian model is available in the Supplementary Materials and Methods (Eq S.6-S.14). As illustrated by Fig 1, the model was designed based on the data generation process of transcriptomics data from iModulons *M* that are differentially activated given conditions *A*. Briefly, the sparse matrix *M* of iModulon parameters captures the sparse relationship between iModulons and genes through the regularised horseshoe prior [16]. Importantly, the membership is defined at the operon level while the particular weight in the iModulon is defined at the gene level. This formulation allowed us to include prior information about the expected operon members per iModulon *p*_0_ (Eq S.7). With this prior, the fitted model characterises membership of operons in iModulons probabilistically. The membership is described, for each operon-iModulon pair, by the distribution of a quantity *κ* (Eq S.8) which lies between 0 and 1: values close to 0 indicate strong membership (true parameter) and close to 1 non-members (zero parameter). Thus, the horseshoe prior establishes the membership decision as a probabilistic inference from the provided data and prior, without any need for post-processing.

**Fig 1.**
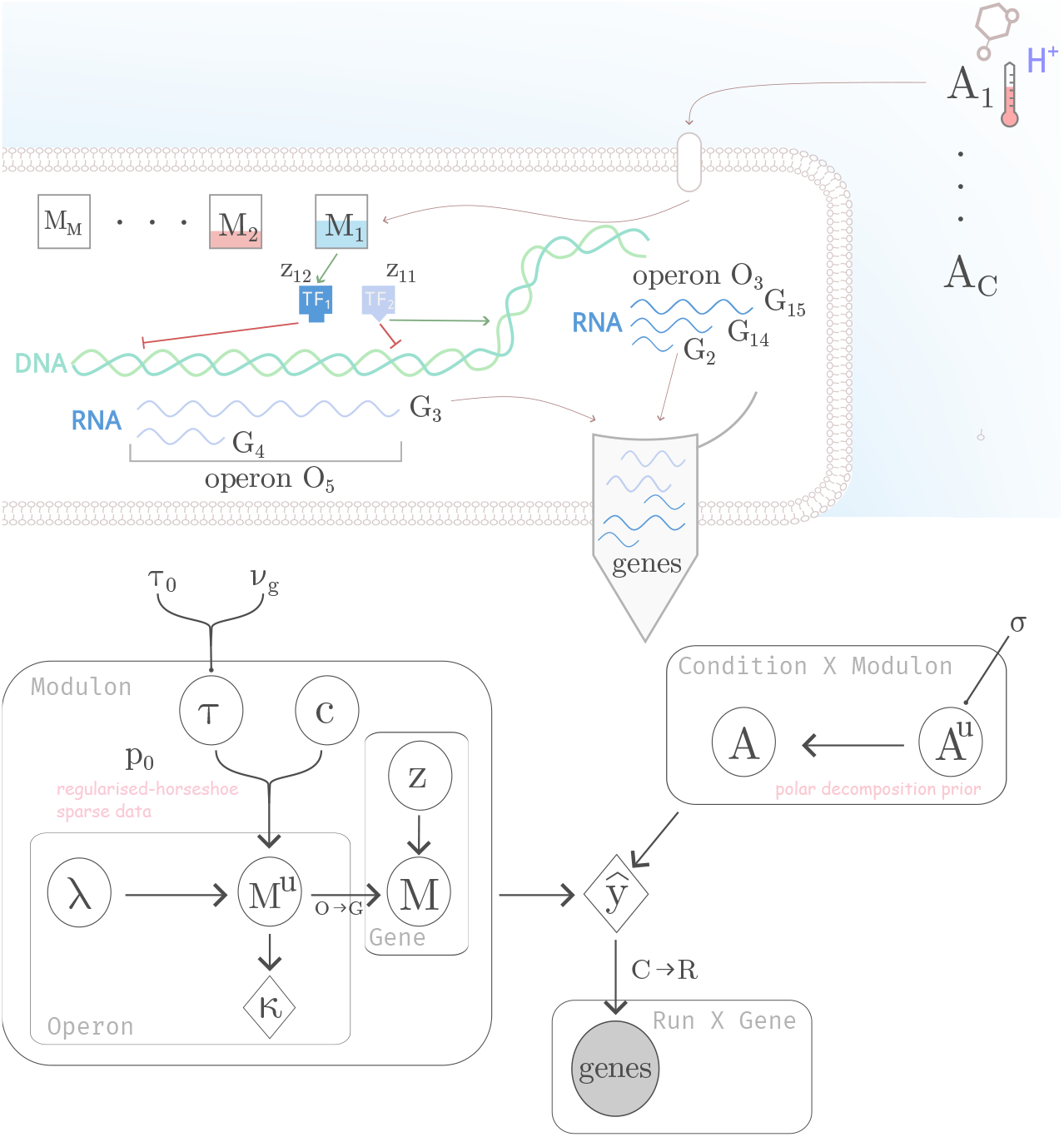
Generative model of transcriptomics data. *(top)* Data generation process of transcriptional regulation. Our model of the mechanism through which cells generate transcriptomics observables. (1) The cell senses the environment in experimental conditions 1, …, *C* and decides how much to activate each of its iModulons, resulting in a set of *M* -vectors *A*_1_, … *A*_*C*_. (2) Given activity vector *A*_1_, each iModulon is activated to a specific extent, illustrated by the colors in boxes *M*_1_, …, *M*_*M*_ . (3) iModulons activate one or more transcription factors, here illustrated by *M*_1_ activating *TF*_1,2_. (4) In turn transcription factors activate or inhibit operons in certain DNA regions (*TF*_1_ inhibits on operon *O*_5_ in the example). (5) Genes in activated operons are transcribed at different expression levels, such as *O*_5_ transcribing *G*3 and *G*4. (6) The mixture of genes is then extracted (gray centrifuge tube) and made available for observation by transcriptomics methods. *(bottom)* Probabilistic graph model of transcriptional response, mirroring the data generation process for transcriptomics. *M*^*u*^ is generated from a regularised horseshoe prior to ensure sparsity, at the operon level, to then be expanded by the individual gene covariates *Z. M* is multiplied by the Activity matrix *A*, orthogonal by construction, to recapitulate the observed log-TPM in the likelihood “genes”. White circles refer to non-observed random variables; gray circles refer to observed variables; rhombuses are deterministic computations and the letters without a shape are hyperparameters.

The activity matrix *A* contains parameters that relate the iModulon activities on each condition. It was made orthogonal through the polar projection [17]) to force orthogonality of the activity of iModulons across conditions. Importantly, only one parameter relates an iModulon to an experimental condition, shared across replicates.

Finally, the likelihood is defined as a linear regression of the matrix product *MA* to the transcriptomics data *y*. When indicated, enforcement of the number of operons per iModulon was also encoded in the likelihood (see Eq S.15).

We tested several variations of this model, including varying the values of hardcoded hyperparameters and altering the model structure, for example, to accommodate data of different nature. These changes are fully detailed in the Supplementary materials and methods.

### Alignment with known transcriptional regulatory network

We evaluated our analysis by recording the alignment of our model’s iModulons with a known TRN from the PRECISE-1. The alignment was performed by exhaustively comparing the genes using the F1-score metric between members of the iModulons and all known single and double TF members (see [7] for details).

The F1-score is the harmonic mean of the precision and recall between two categorical datasets. The F1-score across iModulons gives a goodness-of-fit score indicating how well the iModulons capture biologically relevant information about the set of genes that are known to be co-regulated.

To compare the two methods, we used the clustering method described in [7] to align their results, while the membership function of an operon in a iModulon in the Bayesian generative model is a summary of *κ* (Eq S.8).

The global goodness-of-fit is summarised using the F1-score as in [7], as shown in Eq 1

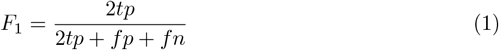

where *tp* refers to the true positives, *fp* is the false positives and *fn*, the false negatives, is the harmonic mean between precision and recall. It is used here to give an estimate of the predictive performance that accounts for the sizes of the overlap between compared sets relative to their sizes.

## Results and discussion

### Bayesian inference of independent modulons of transcription

The Bayesian model presented in the last section and schematized in Fig 1 was fitted using the log-TPM from the PRECISE-1 database [7].

In order to illustrate the properties of the Bayesian iModulons, Fig 2 shows three iModulon—columns of the *GM ⊙ Z* matrix in Eq S.14—with perfect alignment to the regulons chbR+nagC, gadE+gadX and nikR. We note three key characteristics.

**Fig 2.**
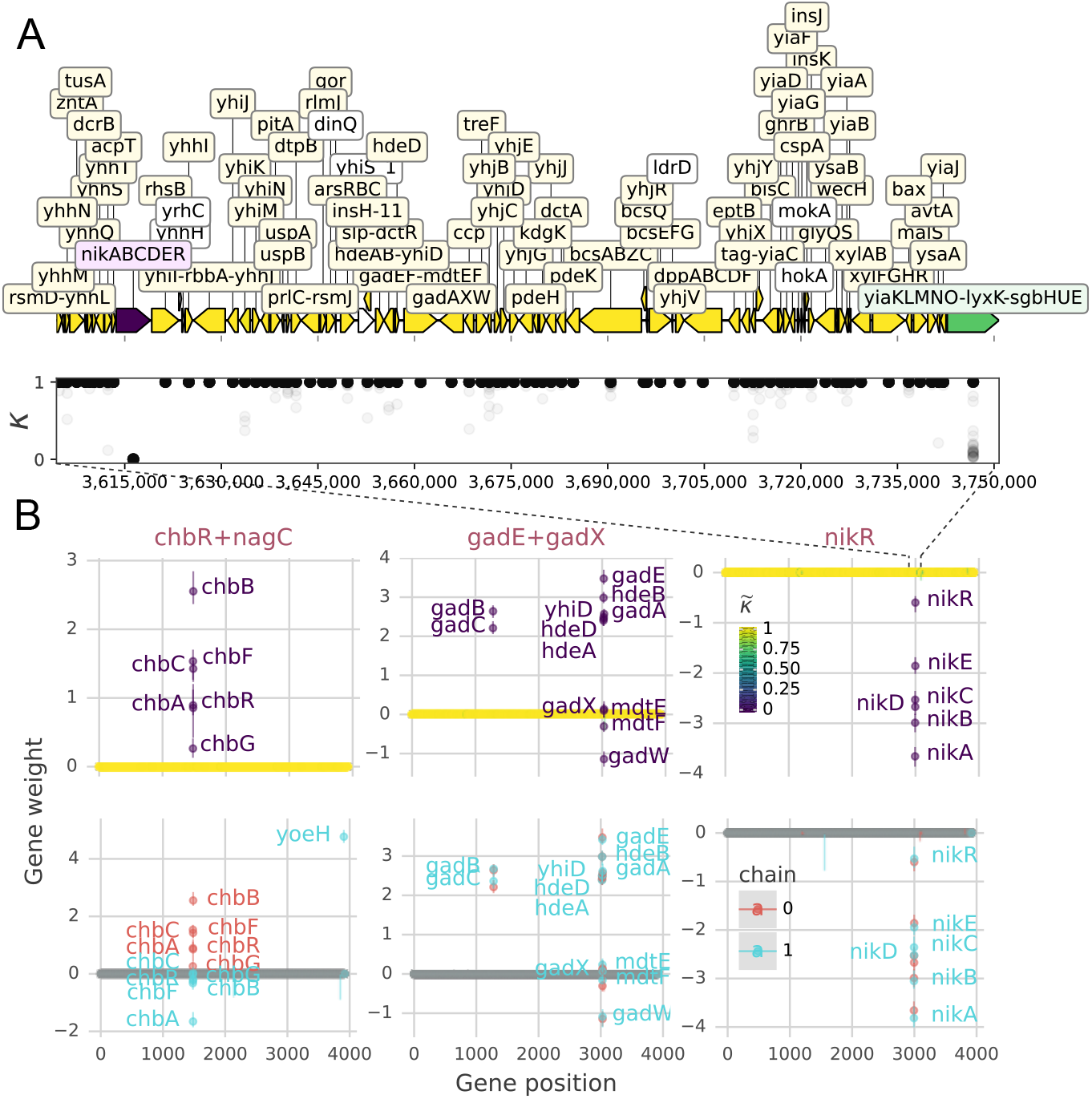
Bayesian sparse iModulons. **A**. *kappa* posterior draws of each operon in the *nikR* iModulon in the genomic vicinity of its only operon member *nikABCDER* plotted with DNA features viewer [18]. The color of the operon, as in B, represents the summary statistic of sparsity 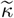 (Eq S.8). **B**. *(top)* Summarised posterior distributions of three iModulons (columns of *GM ⊙ Z* in Eq S.14) that align to the TRN with perfect F1-score, as fitted using Eq S.14 and Eq S.15. *(bottom)* The same iModulons on chains 1 and 2. The points indicate the median value of the *M* posteriors while the line limits are the 25% and 75% percentiles. 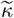 refers to the aggregated sparsity (Eq S.8) of an iModulon parameter, where 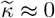 indicates a non-zero parameter; i.e., a gene that is a member of the iModulon. Gene names are shown for values of 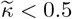.

First, the sparsity of the gene weights. Most posterior weights collapse to zero (Fig 2B). For example, the nikR iModulon (Fig 2A) produces highly skewed *κ* distributions, faithfully captured by the summary statistic 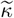 (Eq S.8): it flags both confident members/non-members (purple/yellow) and lower-confidence exclusions such as the *yiaKLMNO* operon (green).

Second, membership is decoupled from gene weight. Because *κ* is independent of the weight *Z*, genes with small weights can still be associated with an iModulon. *GadX* and *mdtE* in the gadE+gadX iModulon illustrate this feature, which is particularly valuable for genes with low expression levels such as transcription factors like GadEWX [1].

Third, credible-interval widths vary by gene and iModulon. For example, the chbR+nagC iModulon shows broader intervals than the other two iModulons in Fig 2, reflecting the higher experimental noise in its RNA-seq profiles.

To quantify goodness-of-fit of the iModulons to the TRN, we calculated the F1-score between each iModulon and TF in the TRN of the RegulonDB as extracted for [7]. The F1-score across iModulons for each method is shown in Table 1.

**Table 1.** F1-score goodness-of-fit percentiles for the alignment with the Transcriptional Regulatory Network.

|  | 75% | 50% | 25% | #MATCHED TFs |
| --- | --- | --- | --- | --- |
| *ICA Baseline* | 0.73 | 0.50 | 0.22 | 60 |
| $Z \in \mathbf{N}_{\mathbf{O}}$ (Eq S.12) | 0.64 (0.03) | 0.52 (0.04) | 0.40 (0.02) | 64.50 (4.93) |
| $Z \in \mathbf{N}_{\mathbf{G}}$ $p_0 = 3$ (Eq S.13) | 0.78 (0.04) | 0.57 (0.02) | 0.45 (0.02) | 68.50 (2.89) |
| $Z \in \mathbf{N}_{\mathbf{G}}$ $p_0 \approx TF$ (Eq S.13) | 0.69 (0.04) | 0.55 (0.01) | 0.42 (0.01) | **70.50 (2.08)** |
| $\iota \sim p_0$ (Eq S.15) | 0.82 (0.05) | **0.67 (0.04)** | 0.50 (0.05) | 59.75 (2.22) |
| $\iota \sim p_0$ (Eq S.15 + S.18) | **0.84 (0.02)** | 0.66 (0.01) | 0.51 (0.03) | 55.00 (3.37) |
| 2D *w/mRNA* (Eq S.15 + S.16) | 0.80 (0.01) | **0.67 (0.03)** | **0.53 (0.03)** | 59.50 (6.03) |
| 2D *w/prot* (Eq S.15 + S.16) | 0.81 (0.05) | 0.60 (0.03) | 0.47 (0.01) | 57.25 (3.59) |

The mean value of percentiles and matched TFs across 4 chains is indicated with the standard deviation in parentheses. “ICA Baseline” corresponds to the result from fitting FastICA with the optimal number of iModulons using the k-means threshold method as described in [7].

Overall, the Bayesian method shows better goodness-of-fit in the 25% percentiles. It can be seen how applying different weights per gene in the iModulon matrix was required to show comparable or better goodness-of-fit than the ICA approach. Moreover, the addition of the effective parameters likelihood in Eq S.15 made the Bayesian model clearly outperform the ICA baseline across percentiles, although the number of matched TFs decreased to a similar level as the baseline.

Penalising the likelihood using Eq S.15 consistently improved alignment at the cost of obtaining fewer TFs. With that in mind, the model that assigns an equal prior of expected operons per iModulon *p*_0_ = 3 (Eq S.13) is still competitive with the ICA Baseline from [7] if information of the TFs were not available, as is expected for most bacteria.

### Global transcriptional shifts in ALE strains explain artificial noise iModulons

A fully identified iModulon with most of the genes in the genomes as members appears across chains and across the runs reported in Table 1. Generally, we would not expect a coordinated mode of regulation that affects nearly the whole genome, therefore we drew two hypotheses for the existence of this “noise iModulon”: that it accounts for basal expression or that it is required to accommodate an outlier condition. We show in S1 Fig that adding an extra basal expression vector did not prevent the appearance of the noise iModulon and that the noise iModulon was associated with one condition in which cells were subjected to Adaptive Laboratory Evolution (ALE). However, removing this condition (`efeU _ _ menFentCubiC_ale38` from [19]) resulted in the appearance of another noise iModulon (data not shown), which was associated with a different ALE condition.

This investigation suggests that the noise iModulon arose from global shifts in some ALE strains, consistent with the fact that ALE often induce mutations in the transcriptional machinery [20–22]. We attempted to address this issue by augmenting our model with a hierarchical component, applying an independent offset to the gene covariates *Z* of the three conditions with highest standard deviations in the Activity matrix (S1 Fig D). As illustrated in Fig 3D, this amounts to assuming that the gene-iModulon membership *M* does not change after ALE, though the particular gene expression may be tuned by ALE mutations.

**Fig 3.**
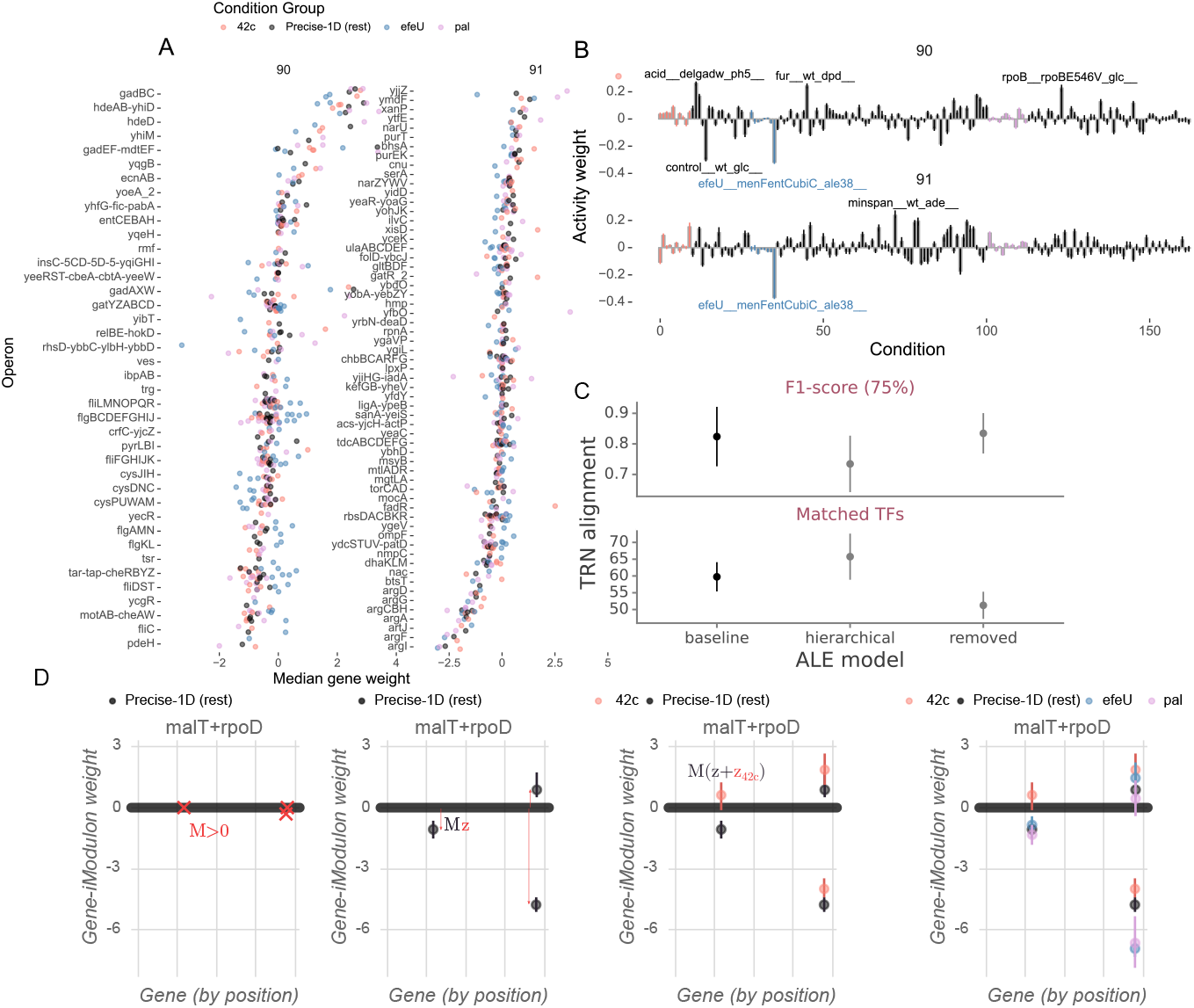
Largest hierarchical iModulons. **A**. Gene weights for the 90 and 91 iModulons in the hierarchical model with different offsets from the normal values for the selected ALE groups, with the operons as labels. **B**. Corresponding activity weights for iModulons 90 and 91. **C**. TRN Alignment performant for the models with the hierarchical link, the normal model and the normal model inferred without the three conditions. The removal of conditions decreases the coverage in 10 matched TFs while the hierarchical model increases coverage with respect to the normal reference model, although sacrificing exact matching of the TRN. **D**. malT+rpoD iModulon separated in its components from the hierarchical prior.

The three experiments with greatest Activity matrix standard deviations in SFig 6D were assigned special offsets for the gene covariate matrix *Z*, as shown in Eq S.19. No noise iModulon was found for this hierarchical model. The two iModulons with the most operon members are shown in Fig 3. iModulon 90 seems to be associated with glutamate-dependent acidic resistance (*gadBC, hdeBC* [23]) and iModulon 91 with Arginine metabolism in the conditions for which no offset was applied. In addition to these genes with large weights in the non-grouped conditions, the weights of genes related to chemotaxis (*tar-tap-cheRBYZ*), motility (*fliL-R, fliC, fliDST, flgB-J, flgAMN, pdeH*) and bacterial competition (*ymdF* [24], *relBE* [25], *rhsD* [26]) were differentially expressed in the ALE strains compared with non-ALE conditions in PRECISE-1. These are all favourable traits for bacterial competition that one would expect in ALE. iModulon 90 is associated with the condition with *gad* deletion, acidic stress and the `efeU_ _menFentCubiC_ale38`. While it might be satisfactory to relate “bacterial competition” with acidic stress and ALE conditions, the gad deletion is hard to explain and might be pointing out to a non-disentangled iModulon. In terms of performance, the hierarchical models showed larger coverage of matched transcription factors to the TRN of *E. coli*, although sacrificing accuracy of exact matches (75 percentile of F1 score) to some extent (Fig 3C).

To sum up, we have demonstrated an approach for detecting outlier conditions and informing the design of subsequent experiments. Furthermore, we showed that hierarchical modelling can separate large “noise” iModulons into biologically meaningful and smaller iModulons. These abilities open the possibility of integrating datasets derived from more diverse sets of strains.

### Smaller (multi-)omics datasets can be included to elucidate new gene regulations

Disentangling iModulons that properly align with biological data requires a large amount of information [7]. Bayesian modelling makes it easier to achieve a sufficient amount of information by allowing diverse data sources to be included at the same time while accommodating their differences.

In this section, different measurement models are added to the initially proposed model in Eq S.14. This addition accounts for the different variability of data collected from different sources, including transcriptomics collected at different times, under different protocols and in different laboratories, while also allowing for the integration of proteomics and transcriptomics measurements in the same model.

The measurement model presented in Eq S.16 was fitted for the compositional transcriptomics (number fractions *ψ*_*m*_) data from [13] in conjunction with the data from PRECISE-1, which was fitted using Eq S.14.

A similar model was used to fit the proteomics data from [12], which corresponds to the transcriptomics later profiled in [13], replacing *ψ*_*m*_ with *ψ*_*p*_. Because the experimental conditions are not identical, the replicate-to-condition mapping (R2) differs from that used for the transcriptomic dataset.

Fig 4 shows three example iModulons inferred using PRECISE-1 and the compositional transcriptomics dataset [13], using *ι ∼ p*_0_ (Eq S.15). It can be seen how new genes (i.e. genes that did not appear in PRECISE-1 because of sequencing depth or different read mappings and assemblies) were added to the iModulons within already established operons (e.g. *trpL*) and as part of different operons not in the PRECISE-1 dataset (e.g. *uof, hokC*). We also fit a model with only the transcriptomics from the 16 conditions from [13] dataset, but this model produced strictly divergent samples (data not shown). The fact that each dataset has genes that do not appear in the other dataset is explained by the fact that [7] used the strict intersection method for feature counts while [13] used the union.

**Fig 4.**
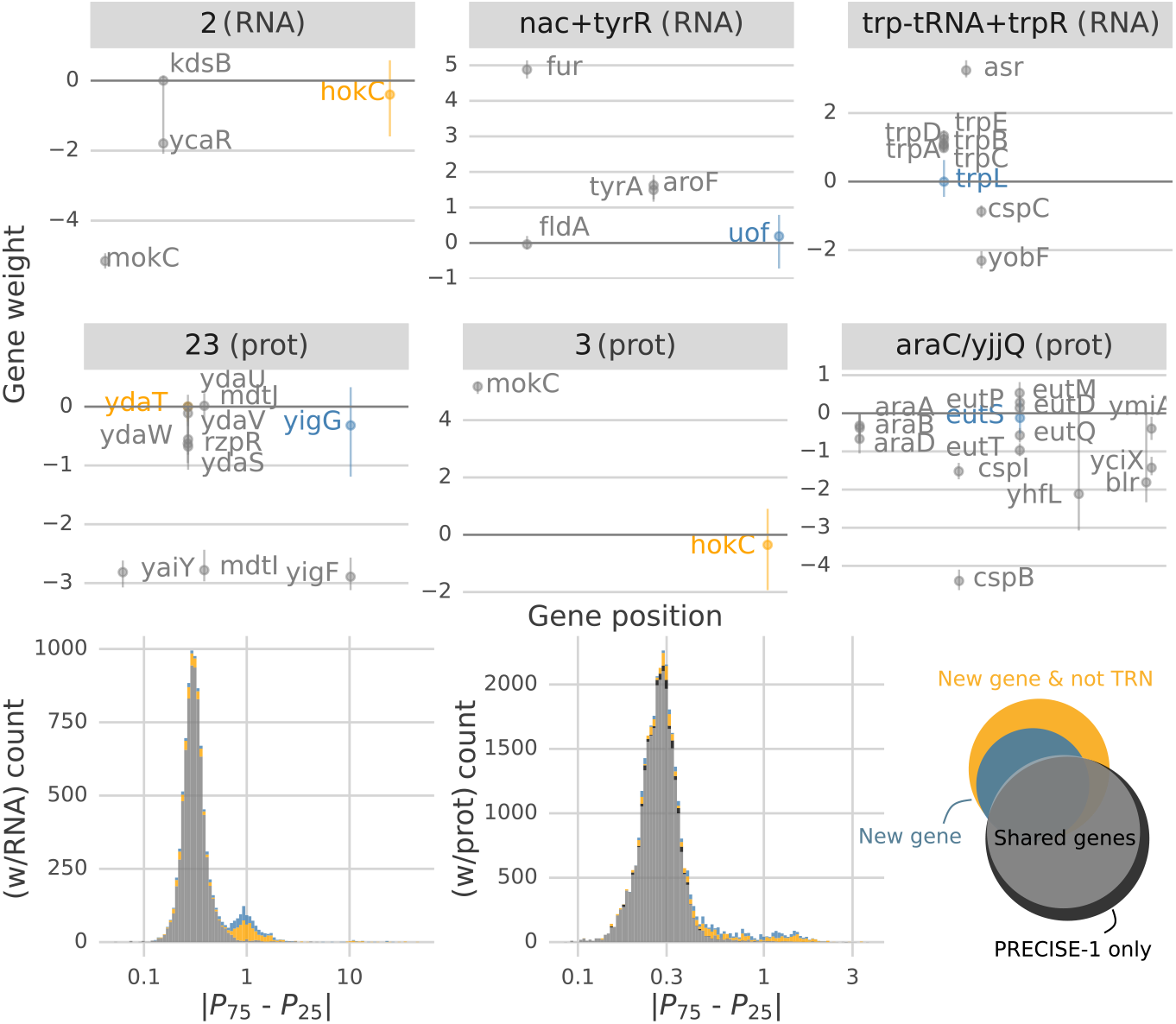
Three iModulons inferred with more than one dataset. Three iModulons (colums of *M* in Eq S.6) from the model with PRECISE-1 [7] and *(top)* the composional transcriptomics [13]; or *(middle)* the compositional proteomics from [12]. *(bottom)* Absolute interquartile difference of gene weights for the models with two datasets: PRECISE-1 with transcriptomics “w/RNA” or proteomics “w/prot”). “New gene” refer to those genes that were in the additional datasets but not in PRECISE-1 and the “New gene & not TRN” refer to those that were also absent in the TRN; the points refer to the median value of the posteriors while the line limits are the 25% and 75% percentile range.

The alignment with the TRN in this case was worse than for the same model with only PRECISE-1: see Table 1. This is unsurprising since, of the 478 genes added from [13], 65.27% are not in the TRN data used for alignment; their inclusion as members can only lower the F1-score. Thus, these results had to be evaluated qualitatively through manual inspection of their biological meaning. Besides the obvious within-operon additions, the added *hokC* in iModulon 2 (Fig 4) is regulated by *mokC* [27]. In the same manner, the open reading frame *uof* [28] in iModulon nac+tyrR is the gene with locus tag `b4637`, which is regulated by *fur* in the same iModulon. The actual genomic position of *uof* —which should be next to *fur* —is misplaced since the operon information was extracted from the same Biocyc table in [7] where that gene was not annotated. This highlights the ability of the method to coalesce information from different datasets to the same latent iModulons in a biologically meaningful way.

Similar examples to those of the transcriptomics dataset are displayed in the middle row of Fig 4, using the proteomics data for the same conditions [12, 13].

The absolute interquartile ranges of iModulon weights for parameters corresponding to genes not in PRECISE-1 (“New genes” in the bottom row of Fig 4) were higher than for other genes. This aligns with the fact that the added datasets have one order of magnitude fewer conditions: 163 conditions in PRECISE-1 compared with 16 in the added datasets. This effect is less pronounced for the added transcriptomics dataset, where the interquartile range is around 1 for both new and non-new genes. However, it should be noted that there was a systematic difference in how gene expression in these two categories was captured in the measurement model, which could also explain the difference. Specifically, genes that were not in PRECISE-1 were affected by an added softmax term before the measurement model as described in Eq S.16.

These results have shown how Bayesian ICA allows for the integration of data from different sources into combined insights by carefully modeling the data generation process of other datasets and omics into a single model.

### A note on invariances between chains

The multiplication of two matrices *A* and *M* in an ICA model produces *permutation* invariances: the positions of columns in *M* or rows in *A* can vary across chains whilst producing the same matrix after multiplication. Additionally, *sign* invariances are produced when neither matrix is constrained positive. This applies to both Bayesian and non-Bayesian ICA.

[7] employed post-processing with sign correction to the maximum value to remove the sign invariance and used clustering to identify “robust” iModulons. The same post-processing could be applied here, based on the quantities *κ*. For the results presented in this work, the chains were aligned using the between chain *κ*_*m*_ F1-score, in the same fashion as the TRN enrichment. This approach is less sophisticated, but we were primarily interested in having a metric of how consistently iModulons agree on gene membership across chains, which could help to compare different prior and likelihood additions regarding the expected iModulon size. With this, we can inspect the raw convergence across chains of Bayesian ICA, while ignoring sign and permutation invariances.

The differences between the chains for *chbR+nagC* in Fig 2 (bottom) illustrate the sign invariance issues of the model. Adding a penalization term (Eq S.15) and applying the post-processing alignment based on the F1-score between chains improved the result: the “nikR” and “gadE+gadX” iModulons show good agreement between chains. However, in the case of the “chbR+nagC” iModulon, the chains disagree about the weights of genes in the *chb* operon, and most importantly, in the membership between the chains in some gene members: gene *yoeH* is included in chain 1 but not in chain 0.

For a global view of the agreement between chains, the F1-score of the iModulons between chains is shown in Fig 5. It can be seen how models with TF-based priors for iModulon sizes show better between-chain membership agreement, leading the F1-score distribution to concentrate on higher values. Interestingly, the likelihood penalization Eq S.15 did not improve membership agreement in comparison with models with information in the prior.

**Fig 5.**
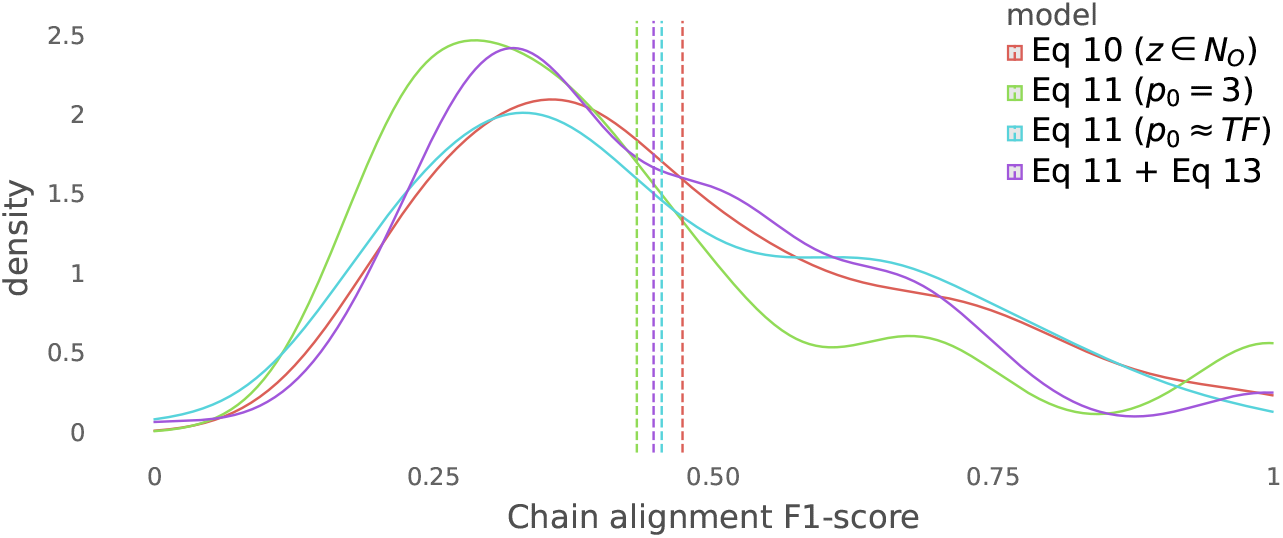
F1-score of alignment of iModulons between chain 1 and 2 of the different models. The dashed lines correspond to the mean of the F1-score of each model.

**Fig 6.**
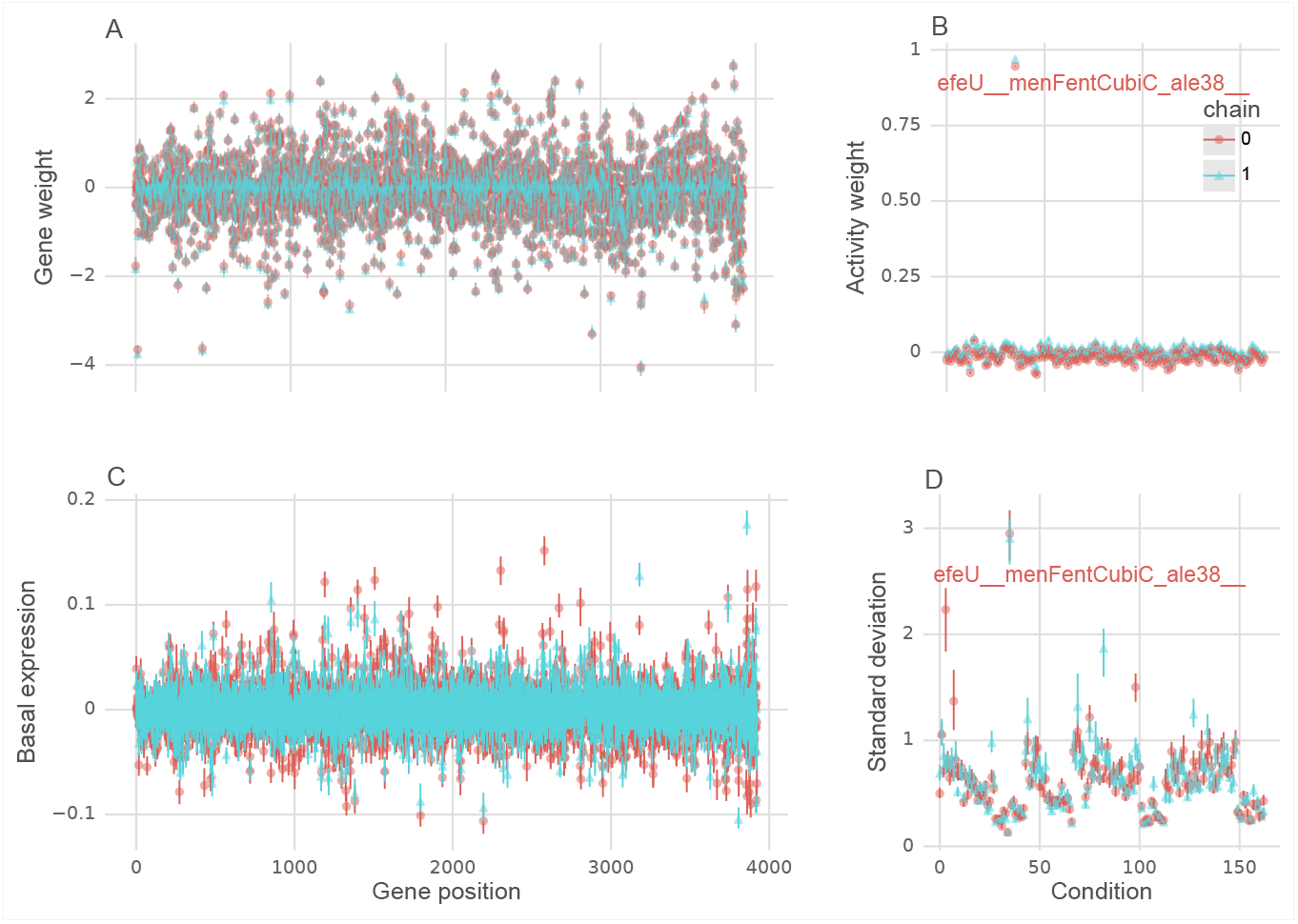
Noise iModulon. *A*. Posterior gene weights of the noise iModulon across chain 1 and 2 of Eq S.14 and Eq S.15. *B*. Activity weights of the same noise iModulon. Only one ALE condition `efeU_ _menFentCubiC _ale38` from [19] appears to be associated with the noise iModulon. *C*. Posterior basal expression per gene on the background vector. The model was modified with a random t-Distributed basal expression vector that multiplies all columns in the activity matrix. Even though the background vector shows some consistency across chains, the characteristic noise iModulon did not disappear from the inferred iModulons. *D*. Per-condition posterior standard deviations of the activity matrix modified by replacing Eq S.10 with Eq S.18, effectively adding a hyperprior to assign different standard deviations per condition in the activity matrix. The points refer to the median value of the posteriors while the line limits are the 25% and 75% percentile range. It can be seen that the highest *σ*_*c*_ posterior associated to an activity was on the same one as the condition associated with the noise iModulon. The posterior on this activity *σ*_*c*_ is identified between chains. This opens the possibility for model-guided design of new conditions to continue on exploring the regulatory landscape of an organism, even for conditions that were not associated with these “noise iModulons”.

## Further work

In this work, the input data has been kept in log-TPM to provide a fair comparison with the normal ICA baseline used in previous approaches. However, it would perhaps be preferable to model the whole data generative model directly from untransformed reads, allowing the model to propagate the uncertainty at a finer level. For instance, the mapping from reads to counts and also the treatment of the counts as generated from a composition instead of log-transforming the data could be included in the generative model. In the same direction, there is a plethora of metadata available in PRECISE-1 (and beyond), such as the base media composition, pH, temperature and sampling time, which could be used as covariates to generate the latent Activity matrix.

More generally, the development of a Bayesian ICA opens for the possibility of including all kinds of information from different sources. Below, we propose different models for researchers with adequate data generation capabilities.

For gene knock-outs we propose the generative model Eq 2,

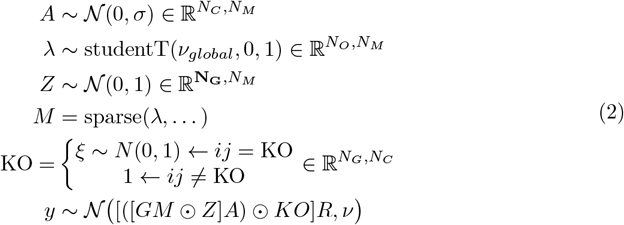

where the information of the knockouts per gene and run is encoded in a matrix KO that adds an inferred effect *ξ* to the gene (or the whole operon where the gene participates) if it was knocked-out in that condition. We hypothesize that this could improve guiding the likelihood, preventing iModulons from being distorted by single gene knock-outs.

As we have demonstrated, Bayesian ICA can be applied to different datasets and different omics data types at the same time, including transcriptomics and proteomics. With a proper alignment of genes based on, for instance, best bidirectional hits, pan-organism iModulons could also be introduced.

The final proposed model is a full translation model using transcriptomics and proteomics for the same conditions, replacing the measurement model in Eq S.14 with Eq 3,

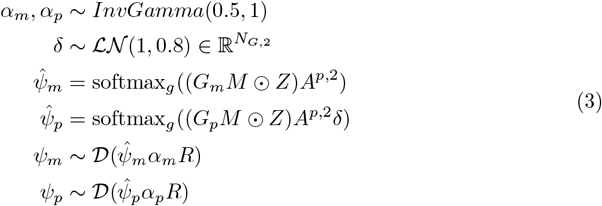

which follows the compositional measurement model in Eq S.16, adding a vector of translation rates *δ* to generate the proteomics from the transcriptomics samples. This involves accounting for the different mappings from the shared operons *M* to the particular gene *G*_*m*_ and protein *G*_*p*_. We hypothesize that, with enough data, this kind of model would calibrate the iModulons more tightly according to their function since the causal transcript-to-protein link would inform the model about the residence time and protein concentration needed to carry out a biological function, while keeping the faster and closer to the signal transcriptomics information.

Other omics data types like metabolomics might be considered for inference. However, as seen in Fig 4, a similar amount of data for different omics types yielded more spread distributions for proteomics than for transcriptomics. Thus, it is not guaranteed that simply adding new omics data types of different variability would be beneficial.

Between-chain agreement remains an area for further improvement. Currently, the recommendation is to only use one chain, as the Bayesian models have demonstrated good alignment with prior biological knowledge. Another shortcoming of the proposed Bayesian model is that, although it provides uncertainty for both the gene membership (in *κ*) and gene weight (in *Z*), it lacks a quantification of the uncertainty for the iModulon as a whole. In other words, we do not yet have a reliable quantitative measure of how certain we can be that an iModulon represents a truly independent and coordinated transcriptional unit. A positive outcome is that all Bayesian models seem to align better with biological regulons in the lower tail (25% percentile, Table 1) than the classical ICA. However, future efforts should focus on finding parameters, maybe in the form of hyperpriors for *M*, that estimate the uncertainty of an iModulon.

## Supporting information Supplementary materials and methods Background

### Independent Component Analysis

Independent Component Analysis assumes that the numbers comprising an *I × J* matrix *X* are weighted sums of a known number *K < J* independent component vectors, as shown in Eq S.4.

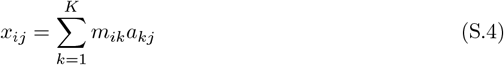

or Eq S.5 in matrix notation

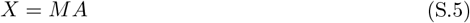

It is assumed that the columns of the matrix *A* are column-wise probabilistically independent, so that the probability of the *j*^*th*^ column of *A* is the product of the *K* marginal probabilities, i.e. 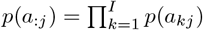. Secondly, it is also assumed that the rows of *A* have non-Gaussian marginal distributions. See [29] for a discussion of an optimisation-based approach to ICA and [30] for a discussion of Bayesian ICA.

In the canonical application of ICA each row of *X* represents a time course of signals from a receiver detecting input from *k* sources, each row of *A* represents the time course of signals from a source and each column of *M* represents how a source mixes between receivers.

ICA can be applied to transcriptomics data analysis by interpreting the observation units as genes rather than receivers, and the observation rows as separate transcriptomics experiments rather than as time courses. Instead of latent sources, the columns of the matrix *M* and rows of the matrix *A* are interpreted as representing “iModulons”, a concept which we explain below.

### iModulons

An iModulon is a hypothetical latent allocation of weights to a subset of genes derived from the results of an analysis involving ICA, that represents a way that a cell could regulate its genes in response to changing conditions.

For example, suppose a certain iModulon I1 regulates just two genes G1 and G2 with respective weights 0.5 and -0.5. When I1 is activated, G1 will be up-regulated and G2 will be down-regulated by the same amount. In contrast, when I1 is deactivated, the opposite regulation will occur. It is typically presumed that there are far fewer iModulons than genes, that an iModulon will substantially up- or down-regulate any gene that it affects, and that most iModulons will affect a small number of genes. For example, the number of genes per iModulon in PRECISE-1K is 17.66 on average with a median of 9 [31].

It is plausible that iModulons roughly describe how gene regulation works because of the known existence of transcription units, transcription factors and regulons. Transcription units are sets of genes that share an RNA binding site, with the result that they can only be regulated jointly. Transcription factors are proteins that activate or deactivate particular transcription units. Regulons are sets of genes that are regulated by exactly the same transcription factors. It is likely that a latent representation like iModulons is roughly correct, in particular, based on the high degree of overlap between iModulons characterised in *Escherichia coli* and the experimentally derived regulons from regulonDB [7], and it is clear that most transcription factors affect a small number of genes and that there are fewer transcription factors than genes. Nonetheless, iModulons are not perfectly mappable to transcription factors, as they often correspond to the combined effect of more than one transcription factor or to different modes of the same transcription factor, such as *Crp* in PRECISE-1 [7].

### Bayesian generative model

We constructed a new Bayesian generative model implementing Independent Component Analysis, using the model presented in [30] as a starting point. Like the model in [30], our model treats the values of the ICA *M* and *A* matrices as unknown parameters, ensures that the conditions for ICA are met via the choice of prior distributions for these parameters and represents measurements using a regression model that connects the ICA *X* matrix with the observed transcriptomics data.

The main differences between our modelling approach and the one in [30] are as follows:

- We fit our model using gradient-based MCMC sampling, whereas [30] performed approximate Bayesian inference via expectation-maximisation.
- In our analysis each value of the ICA mixing matrix *M* is unconstrained, and can in principle be any real number, whereas [30] imposed a positivity constraint on these values.
- We use a sparsity-inducing prior for the values of *M* .
- We use a normal linear regression whereas [30] use a mixture of Gaussians to describe measurements.
- Our signal matrix (ICA activity matrix *A*) is constrained to be orthogonal.

The following sections elaborate on these differences.

### Sparsity-incucing prior for the ICA *M* matrix

Our model assigns the columns of the ICA mixing matrix *M* independent sparsity-inducing prior distributions, implementing the regularised horseshoe prior presented in [16]. This formulation is shown in Eq S.6. We parametrise *M* using two matrices of auxiliary parameters with the same dimensions, which we call *λ* and *Z*, as well as parameter vectors *c* and *τ* with the dimension of the number of iModulons, using the following formulation:

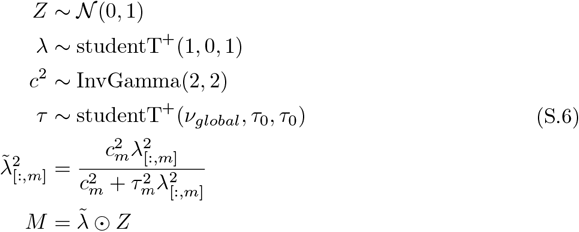

where studentT is the Student’s t-distribution, *InvGamma* is the inverse Gamma distribution; the superscript ^+^, as in *studentT* ^+^, indicates that a distribution is truncated below at zero; the terms *λ*_[:,*m*]_ and 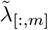 refer to the *m*-th columns of the matrices *λ* and 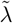; and the symbol *⊙* denotes the Hadamard product, i.e., element-wise multiplication. For more stable sampling we parametrised the student T distribution as a mixture of Gamma distributions in this case, as recommended by [32].

This choice of prior distribution allows our model to include both information about the likely sizes of non-zero values (encoded in the prior for the parameter vector *c*), leading to improved fitting via regularization, and also information about the likely number of non-zero values per column, representing gene members in the iModulons. Eq S.7 shows how this can be done in practice, following the recommendations in [16]. Specifically, it shows how to use information in the form of the expected number of gene members for an iModulon, *p*_0_, to determine the quantity *τ*_0_, which is the location parameter of the prior for *τ* in Eq S.6.

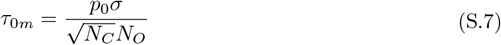

In Eq S.7, *N*_*C*_ is the number of conditions and *N*_*O*_ is the number of operons; *p*_0_ is the expected number of operon members per iModulon (a scalar or a vector per iModulon); and *σ* indicates the *noise level* of the *τ* prior.

For our application, we followed two strategies for exploiting information about the number of gene members in the iModulons: we either used a *p*_0_ of expected genes per iModulon, interpolated from the *in vivo* promoter response data from [33] down to the number of iModulons, or else, when indicated, set the expected number of gene members to 3 for all iModulons.

An important output of the regularised horseshoe prior is the characteristic sparsity quantity *κ*, which in our application indicates whether or not a gene is a member of an iModulon. The *κ* quantity *κ*_*m*_ for iModulon *m* is defined as in Eq S.8 (Eq 2.4 of [16])

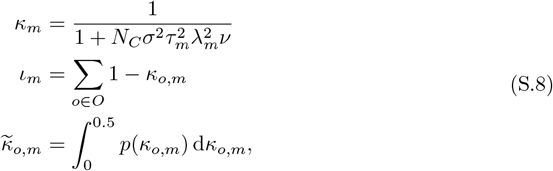

where *ν*is the measurement noise; *ι*_*m*_ indicates the number of effective operons in modulon *m*; *κ* is a distribution in [0, 1] where 0 indicates membership of the operon *o* in the modulon *m*. The aggregation of the *κ* distribution for each gene and operon 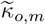, effectively the cumulative distribution function up to 0.5, indicates the proportion of posterior samples that are evaluated as members of the modulon *m*. 0.5 is chosen since it is the “bottom of the horseshoe”: lower values represent membership and higher values non-membership.

For simplicity, we denote the transformation that yields *M* from the auxiliary parameters *λ, τ* and *c* in Eq S.6 as “sparse” in the manuscript formulas, as in Eq S.9, separated from the *Z* for convenience of manipulation as we will see below.

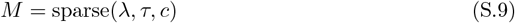

### Orthogonal activity matrix

To ensure that the the ICA activity matrix *A* is orthogonal, we derive it by transforming a matrix *A*^*u*^ of auxiliary variables with the same dimensions. The values of *A*^*u*^ have independent half-normal prior distributions with location parameter 0 and scale parameter 0.5, as shown in Eq S.10.

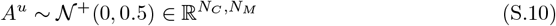

As with optimization-based ICA, our model assumes that the data were whitened and standardised, so that standard deviation 0.5 represents approximately half a standard deviation relative to the overall variation in RNA expression in the PRECISE-1 database.

To generate an orthogonal matrix *A*, we then apply the polar projection from [17], as shown in Eq S.11

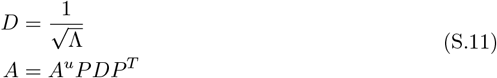

where *P* and Λ are eigenvectors and eigenvalues of the *A*^*u*^ cross-product (see S1 Appendix for a proof that explains this transformation). [17] showed that this transformation is amenable to Monte Carlo methods like Hamiltonian Monte Carlo but produces a rotational invariance.

### Measurement model

The measurement model is a simple linear regression model with a normal likelihood with known measurement error *ν*over the generated operon expression data that arises from multiplying *M* and *A*. In the hypothetical case where operon activity could be measured directly this model would be as shown in Eq S.12:

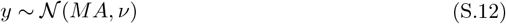

However, we aimed to extract information from measurements of gene, rather than operon activity, taking into account that the expression levels of genes within an operon can differ. Furthermore, our data contained biological replicates.

To accommodate these complications, we first introduced two mapping matrices: *G* with dimensions *N*_*G*_ times *N*_*O*_, i.e. number of genes times number of operons, and *R* with dimensions *N*_*c*_ times *N*_*r*_, i.e. number of biologically unique conditions times total number of possibly-replicated conditions. Every value *G*_[*g*,*o*]_ is 1 if operon *o* contains gene *g* and 0 otherwise; similarly *R*_[*c*,*r*]_ is 1 if condition *c* corresponds to possible replicate *r*. The matrix *GM* then contains the operon activity for each gene in each iModulon, according to *M*, and the matrix *AR* contains the iModulon activation level in each condition, according to matrix *A* whose columns represent only biologically unique conditions.

To allow for differing expression levels for genes within the same operon, we offset the values of *GM* by a nuisance parameter matrix *Z*, with dimensions *N*_*G*_ times *N*_*M*_ and a standard normal prior, with the final measurement model as shown in Eq S.13:

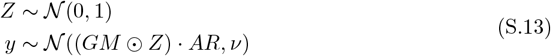

In our case, we set the value of *ν*to 0.3, which is approximately 2 times the 75th percentile of the standard deviation of the same-gene expression between biological replicates. The operon information for *Escherichia coli* was extracted from Biocyc [34].

The full generative model is shown in Eq S.14 for reference.

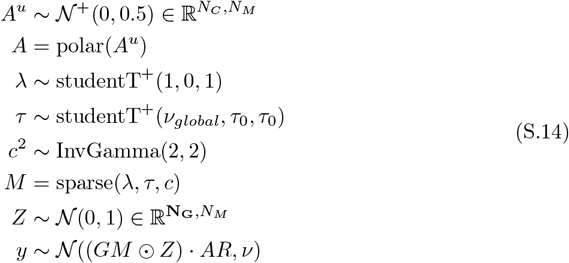

where “polar” denotes Eq S.11 and “sparse” denotes Eq S.9.

### Permutation and sign invariances

Permutation and sign invariances are produced when the two matrices in Eq S.13 are multiplied, which is a challenge for identifiability between chains. The permutation invariance is due to the fact that the same iModulon can take any column position arbitrarily in *M* to then be multiplied by a row in *A*. To address this issue, we impose a likelihood over *ι* in Eq S.8 to *p*_0_ (expected number of members) to ensure that the iModulons are ordered by the number of operons members as described in Eq S.15.

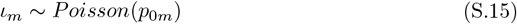

This likelihood term is in addition to the prior in the size of the iModulons in Eq S.7 since this prior only ensures that the iModulons are around the same order of magnitude (as noted in [16] and empirically verified in SFig 7).

**Fig 7.**
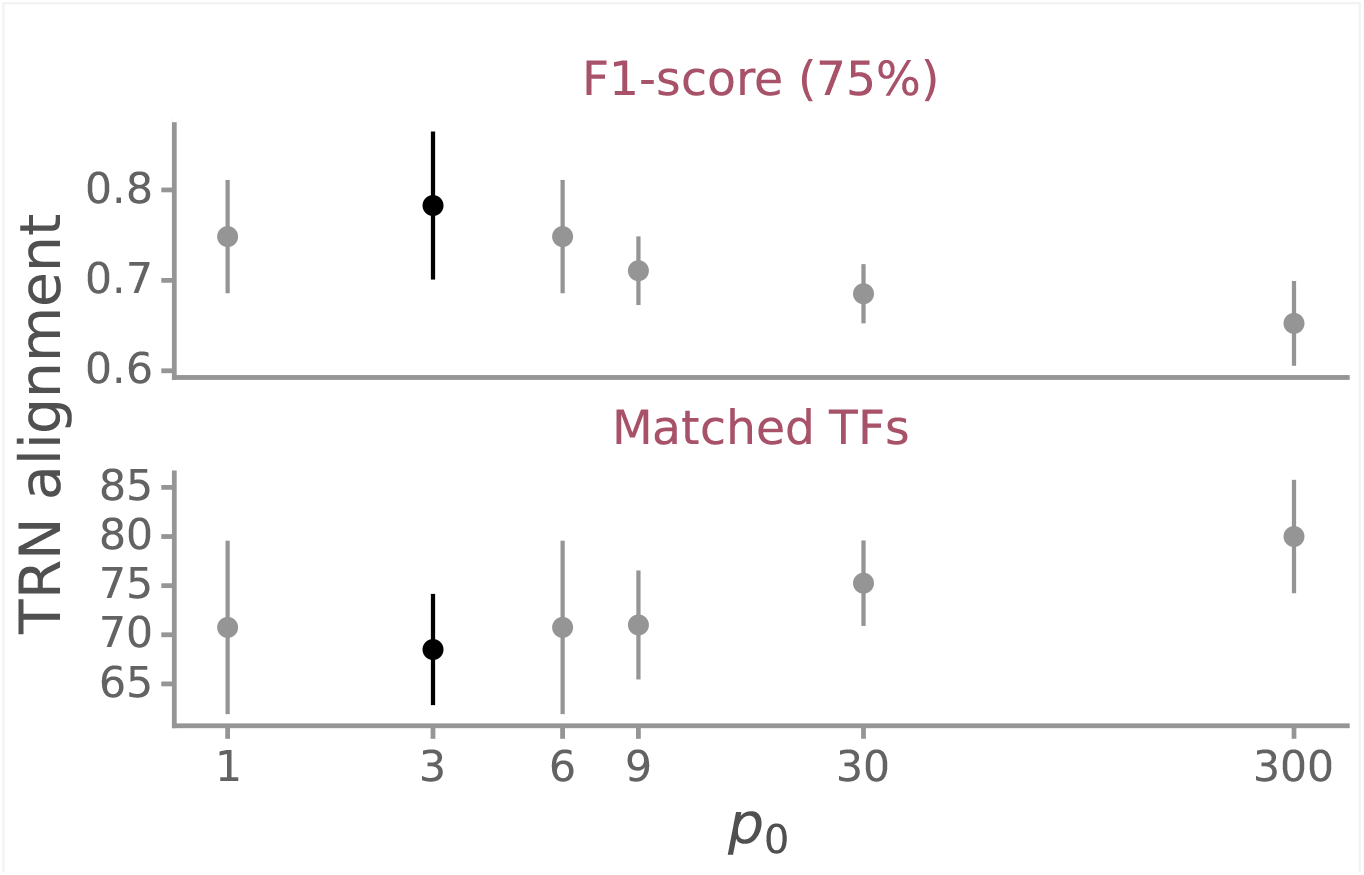
F1-score 75 percentiles and matched TFs across chains for varying numbers of the fixed hyperparameter *p*_0_. The dots refer to the mean of and the error bars to the 95% interval across chains.

### Measurement models for compositional data

The transcriptomics data from [13] contains number fractions *ψ*_*m*_ instead of log-TPM.

Since the number fractions are compositional vectors—i.e. positive-constrained and closed to 1—this kind of data requires a different measurement model. We modelled these data jointly with the PRECISE-1 dataset, using the Dirichlet regression model shown in Eq S.16 for the compositional measurements:

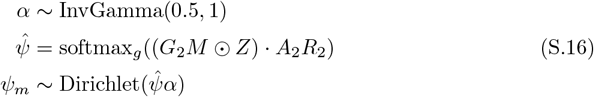

where *ψ*_*m*_ are the observed number fractions in the data, Dirichlet is the Dirichlet distribution applied row-wise, and the 2 indices refer to this second compositional dataset. Notice how the genes in the first dataset are overlapping but not the same as in the second and, hence, they have to be indexed by a new operon-to-gene matrix *G*_2_ to correctly map to the genes in *ψ*_*m*_. The *α* parameter is introduced to allow the dispersion of the members in the Dirichlet distribution to be fitted during inference.

The softmax (Eq S.17) function applied to each gene makes the rows stochastic vectors, as expected in the data:

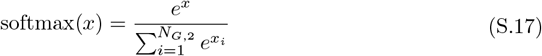

A second model with the same structure was used to fit the proteomics data from [13], replacing *ψ*_*m*_ with *ψ*_*p*_. It is important to note that the conditions in the proteomics dataset are not exactly the same and, as a result, the replicate matrix *R*_2_ is different in this case.

### Condition-specific activity standard deviation

To quantify the uncertainty per-condition, Eq S.10 was substituted with Eq S.18 when finding the prior distribution for a column 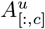 of the auxiliary activity matrix *A*^*u*^:

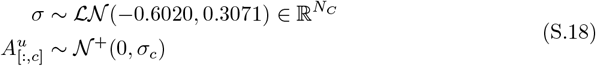

where *σ* is a vector whose length is the number of conditions, with the location and scale parameters -0.6020 and 0.3071 chosen to center around 0.5, with 95% of the total probability mass lying between 0.3 and 1. 0.5 was found to be the value that produces the best TRN alignment for the PRECISE-1 database.

### Hierarchical model of multi-strain gene weights *Z*

To model possible structural differences between groups of conditions, we modified the model in Eq S.14 using a hierarchical model for the parameter matrix *Z* and the consequent changes in the measurement model as described in Eq S.19

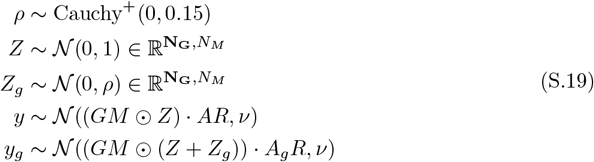

where *y*_*g*_ and *A*_*g*_ indicate the measurement and iModulon activities belonging to a particular group: in our case the ALE conditions (“42c” [19], “pal” [35] and “efeU” [36]). For other conditions, the model is as in Eq S.14. Notice that the *M* matrix is shared across groups, such that the iModulons remain the same in members, but are offset differently.

## S1 Appendix. Polar projection proof

Given a random matrix *X*, we take the eigendecomposition of its crossproduct:

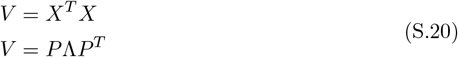

Notice that *V* has unique eigenvalues Λ since it is a symmetric matrix and, thus, orthogonal eigenvectors *P* . Hence, we define the orthogonal matrix *A*

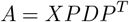

which is orthogonal iff *A*^*T*^ *A* = *I*;

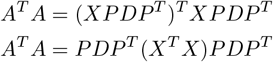

substitute *X*^*T*^ *X* by Eq S.20;

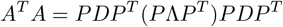

apply *P*^*T*^ *P* = *I* since the eigenvectors are orthogonal;

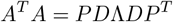

Eq S.11 is designed such that *D*Λ*D* = *I*;

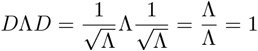

and thus,

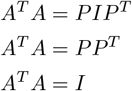

**S1 Fig. Noise iModulon investigation**. The figure shows how the generative model was expanded to test hypotheses about the existence of a big iModulon encompassing a larget set of the genes in *Escherichia coli*.

**S2 Fig. p0 sensitivity**. F1-score 75 percentile and matched TFs summarised across chains for models without *κ* penalty (Eq S.15) and fixed *p*_0_ to different values. It can be seen how the results change only between orders of magnitude, decreasing accuracy to perfect matches (75 percentile of F1-score) when the parameter is change is made larger. Since the resulting iModulons contain more members with larger *p*_0_, the number of matched TFs is also larger.

## Acknowledgments

This work was supported by the Novo Nordisk Foundation [NNF Grant Numbers: NNF20CC0035580, NNF14OC0009473].

## Notes

### Competing Interest Statement

The authors have declared no competing interest.

